# Neuronal modulation of the superior colliculus associated with visual spatial attention represents perceptual sensitivity, independent of perceptual decision and motor biases

**DOI:** 10.1101/2025.04.01.646682

**Authors:** Supriya Ghosh, John H.R. Maunsell

## Abstract

Neurons in the superior colliculus (SC), like those in the cerebral cortex, are strongly modulated in response to shifts in attention, but make a contribution that is distinct from attention-related modulations in visual cortex. It has been a point of contention whether attention-related enhancement of neuronal activity in SC is associated with selective increase in behavioral sensitivity (*d’)* in neurons’ response fields or with animals’ decision bias, which is closely linked with motor planning. By independently controlling monkeys’ perceptual decision and motor criterion, we show that SC activity is strongly correlated with perceptual sensitivity at the neuron’s response field. Responses of the same SC neurons were unchanged in the face of correspondingly large changes in the perceptual decision criterion. Furthermore, the SC activity did not convey information about perceptual detection on individual trials. These results suggest that the SC contributes to the component of attentional states related to heightened perceptual sensitivity.

## INTRODUCTION

Perceptual performance greatly improves with selective attention. There are many distinct perceptual, decision and motor related factors that can independently alter attentional performance ^1,2^. Neuronal activity in numerous cortical and subcortical brain regions changes with shifts in attention ^3-6^. Previous studies examining attention-related modulations in the context of signal detection theory ^7^ suggest that neuronal modulation in visual cortical area V4 is associate with only perceptual sensitivity, or *d’*, and not with the decision criterion ^8,9^. The superior colliculus (SC), a midbrain subcortical structure, plays a crucial role in visual perceptual performance ^5,10-12^. Moreover, it is believed that the SC contributes to visual attention through mechanisms that are distinct from those in the visual cortex ^13^. Responses of sensory-motor neurons in the SC are known to be strongly modulated when animals enhance their perceptual *d’* at the neuron’s response field (RF) ^5,14^. It has alternatively been proposed that the primary contribution of the SC to attentional performance is related to perceptual decision criteria or thresholds ^11,12,15^. The precise mechanisms underlying perceptual behavior mediated by SC neuronal responses have been subject to ongoing debate.

Different neuronal groups within the SC exhibit distinct response properties and varying degrees of spike modulation from sensory, cognitive and motor signals ^16^. Neurons located in the intermediate and deep layers of the SC are more sensitive to various cognitive states compared to sensory neurons in the superficial layer of the SC. Previous studies suggest that the contribution of SC neurons to perceptual performance is largely attributable to sensory-motor neurons in perceptual detection during visually guided behavior ^14,15,17^. However, the relative contributions of these SC neurons to the distinct perceptual and behavioral components of attentional performance remain poorly understood.

To address this, we recorded spikes from multiple neurons in the SC of rhesus monkeys while they performed a novel visual orientation-change detection task ^18^ that cleanly dissociates perceptual *d’* from decision criteria and motor planning. The results revealed that SC neurons increase their firing rate with increased in attention (selective behavioral *d’*) to their RF location. This attentional modulation was more pronounced for visuomotor neurons compared to purely visual neurons, and was virtually absent in motor neurons. Unlike selective behavioral *d’*, changes in perceptual decision criteria had no impact on the spiking of these same SC neurons. These results offer a comprehensive account of how distinct sensory and cognitive components relevant to behavioral performance may be represented in the SC. This behavioral task holds particular significance in isolating specific behavioral components associated with attention states in brain areas that respond to a multitude of signals.

## RESULTS

### Spike modulation of SC visuo-motor neurons associated with changes in response criterion

We independently controlled the motor response criterion between low and high levels while maintaining uniform perceptual *d’* in two rhesus monkeys using a visual orientation change detection task ^18^ (**Fig. 1a**; **Methods**). The animal fixated at a central spot (0.2° diameter) at the start of each trial, and maintained gaze on this spot until it went off. After a variable time of fixation (400-700 ms) two Gabor sample stimuli appeared for 200 ms, one in each hemifield. Following a brief delay (200-300 ms), a single Gabor test stimulus appeared for 200 ms in one of the two locations with a 50% probability of having same or a different orientation than the sample stimulus that had appeared in that location. The monkey had to report whether the test Gabor orientation was a match (0°) or non-match to the preceding sample Gabor (median thresholds (interquartile range), monkey S, 27° (25°-40°); monkey P, 45° (38°-50°)) by making a saccade to the appropriate saccade target after the fixation spot went off (go cue) to receive a juice reward for correct detection. Match and non-match saccade targets (different color and shape) appeared 100-160 ms after the test stimulus was turned off, in opposite locations orthogonal to the sample stimuli. The locations of match and non-match saccade targets remained fixed within a trial block and were interchanged over interleaved trial blocks. The block-wise predictability of the saccade target locations made it possible to measure and control animals’ motor intention or motor response criteria.

**Figure 1.**
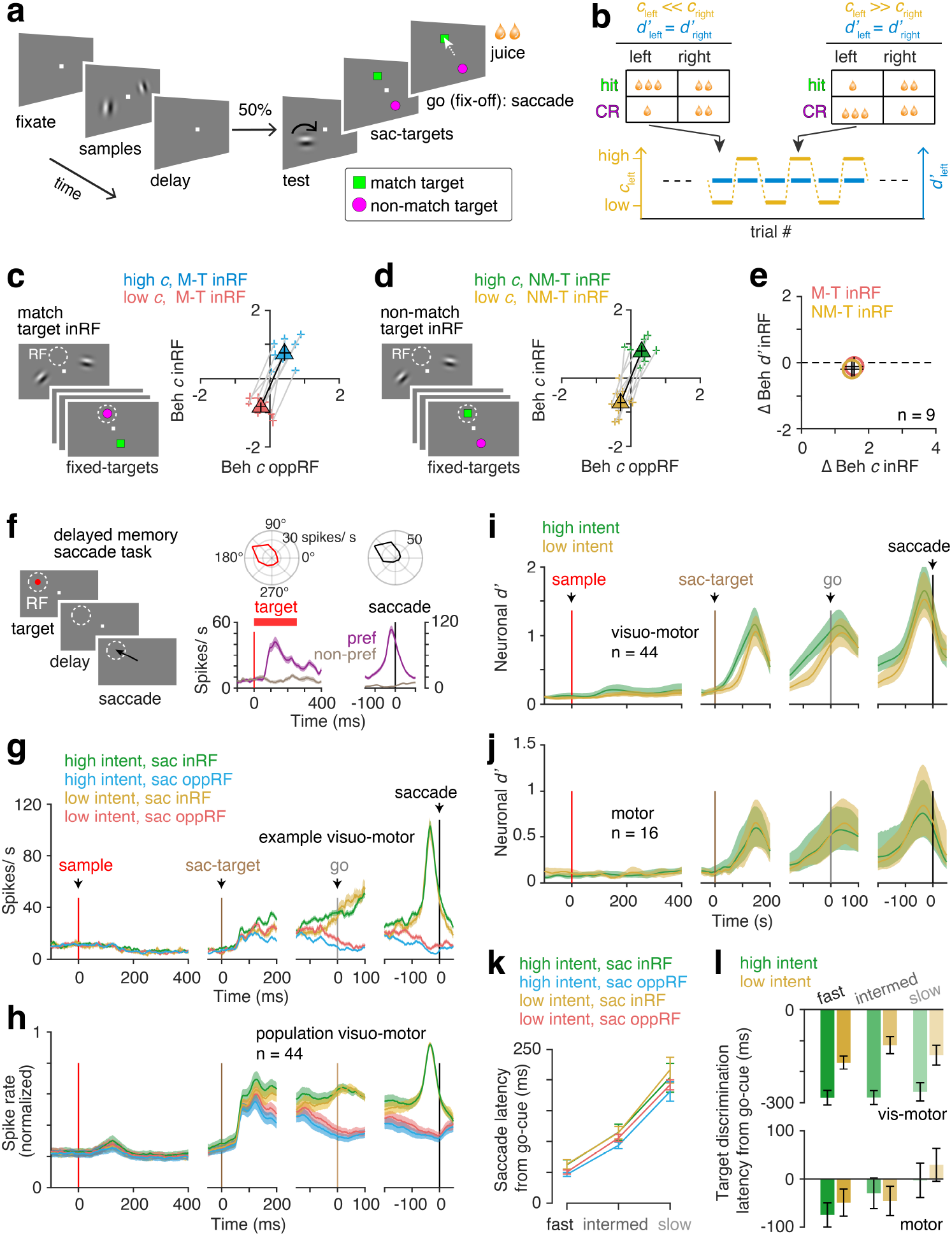
Spike modulation of SC neurons associated with response criterion. (**a**) Visual orientation change detection task (**Methods**). Monkeys fixated (400–700 ms) while attending to one of two sample Gabor stimuli (200 ms) in opposite hemifields. After a short delay (200–300 ms), a test stimulus (200 ms) appeared at one of the stimulus locations. Monkeys reported orientation changes between the sample and the test stimuli by making a saccade to the correct target (distinguished by color and shape) for a juice reward once the fixation-spot went off (go-cue). Saccade target positions remained fixed withing a trial block. (**b**) Monkeys’ behavioral response criterion (*c*) for a stimulus was controlled between low and high across interleaved trial blocks, while behavioral sensitivity (*d’*) at the two stimulus locations remained constant by adjusting reward size for correct detections (**Methods**). (**c-d**) Monkeys’ response *c* across sessions (monkey P, 4; monkey S, 5) for low and high response *c* conditions when the match saccade target was at the neuron’s RF location (c) or in the opposite hemifield (d). Gray lines, individual sessions. Filled triangles, session averages. Error bars, 95% confidence intervals. (**e**) Average changes in behavioral *c* and *d’* in same experimental sessions. While *d’* remained stable, *c* varied between low and high values. Error bars, 95% confidence intervals. (**f**) *Left*, Delayed memory saccade task for classifying SC neurons into visuo-motor, visual, and motor neurons (**Methods**). *Top right*, visual and saccade-related spike responses of an example SC visual-motor neuron. *Bottom right*, PSTHs aligned to visual target and saccade onsets at the preferred and non-preferred RFs. Error bars, ± 1 SEM. (**g**) PSTHs for the same example visuo-motor SC neuron in (f), grouped by saccade location and response bias level (saccade target selection). Spike rates are aligned to onsets of sample stimuli, saccade targets, go cue and saccade. Error bars, ± 1 SEM. (**h**) Same as (g) for population visuo-motor neurons (n = 44). (**i-j**) Neuronal *d’* for SC visuo-motor (i) and motor (j) neurons for selecting a saccade target in the RF for different levels of response biases. (**k**) Mean (± 1 SEM) saccade latencies for different response biases and saccade target selections. Saccade latency from the go cue was grouped into three quantiles. (**l**) Neuronal discrimination latency (*top*, visuo-motor neuron; *bottom*, motor neurons) for selecting saccade targets at the RF versus opposite location, grouped by response bias levels and saccade latency as in (k) (**Methods**).

To control response criteria, monkeys were instructed to distribute their spatial attention equally to the two locations while shifting their response criteria to the two locations in interleaved small blocks of trials (monkey P, 120-140 trials, monkey S, 120-160 trials) (**Fig. 1b**). Criteria was switched between low and high values at one location independent of behavioral *d’* by adjusting the reward size ratio for hits and correct rejections (CR) at that location while keeping average reward sizes the same between the two locations.

Monkeys selectively shifted their behavioral criteria between low and high values when the match saccade target was in the RF (F_(1, 16)_ = 14.96, p < 10^−7^, repeated measures ANOVA, **Fig. 1c**), as well as when the non-match saccade target was in the RF (F_(1, 16)_ = 15.10, p < 10^−7^, **Fig. 1d**), without any changes in behavioral *d’* (match saccade target in-RF, F_(1, 16)_ = 0.02, p = 0.84; non-match saccade target in-RF, F_(1, 16)_ = 0.08, p = 0.58). This independent control of behavioral criteria and perceptual sensitivity (**Fig. 1e**) was crucial for isolating and investigating the neuronal modulations of SC neurons associated with response bias in attentional performance.

### Spike modulation of SC neurons associated with changes in response bias or motor intent

We recorded spiking activities from populations of SC neurons while the monkeys’ response criteria was controlled while behavioral *d’* remained unchanged during the visual attention demanding task (monkey S, n = 36; monkey P, n = 33). SC neurons were classified as visuo-motor, visual and motor neurons based on the selectivity of visual and saccadic responses using a delayed memory visual task (visuo-motor, n = 44; visual, n = 9; motor, n = 16 from both animals; **Fig. 1f & Supplementary Fig. S1**; **Methods**).

Visuo-motor neurons exhibited increased spiking to the saccade target stimuli during trial blocks when the monkey’s response bias at the RF was high (**Supplementary Fig. S1**). This occurred either when the behavioral criterion was low and the non-match saccade target was at the RF, or when the behavioral criterion was high and the match saccade target was at the RF. Neuronal spike modulation associated with changes in behavioral criteria (Δ*c*) was measured using neuronal *d’* based on the mean spike counts over 200 ms (50 to 250 ms from saccade target onset, **Methods**). There was a significant spike modulation as measured by neuronal *d’* for behavioral Δ*c*, in visuo-motor neurons (neuronal *d’*, for nonmatch saccade target in-RF, mean ± SEM = 1.03 ± 0.15; *P* < 10^−6^; for match saccade target in-RF, mean ± SEM = 1.07 ± 0.12; *P* < 10^−7^, signed rank test). There was no neuronal modulation in SC visual neurons for behavioral Δ*c* (neuronal *d’*, for nonmatch saccade target in-RF, mean ± SEM = 0.17 ± 0.36; *P* = 0.98; for match saccade target in-RF, mean ± SEM = 0.28 ± 0.52; *P* = 0.2, signed rank test). Response modulation of SC motor neurons with behavioral Δ*c* were weak, but significant (for nonmatch saccade target in-RF, mean ± SEM = 0.67 ± 0.27 *P* = 0.03; for match saccade target in-RF, mean ± SEM = 0.88 ± 0.34; *P* = 0.03, signed rank test).

In order to dissociate the spike modulation associated with the response bias (also referred to as ‘motor intent’, or ‘motor planning’) from the physical motor execution, trials were classified according to the monkeys’ saccade location (in-RF or opposite-RF) and the level of motor intent (high versus low response bias) in selecting a saccade location. Spike rates of visuo-motor neurons following the onset of saccade targets increased with a higher level of the monkeys’ intent to select a saccade target within the RF compared to the opposite location (*green* versus *yellow* traces, **Figs. 1g-h**). Neuronal *d’* increased significantly after the onset of saccade targets with higher motor intent (**Fig. 1i**).

In contrast, motor and visual neurons showed no modulation in response to changes in motor intent (**Figs. 1j & Supplementary Fig. S2**). We further looked at whether the motor intent-related neuronal modulation reflected the latency of saccade initiation. All trials were grouped into fast, intermediate, and slow quantiles based on the saccade latency relative to the go-cue, separately for each trial type. (**Fig. 1k**). The neuronal latency of saccade target selection for individual visuo-motor and motor neurons was defined as the earliest time, relative to the go-cue, when decoding accuracy (in-RF vs. opposite-RF) was significantly higher (p < 0.05 over at least 80 ms) than the shuffled trials. This was assessed for each set of motor intents and saccade latencies. (**Fig. 1l; Methods**). The neuronal latency of saccade target selection in visuo-motor neurons was significantly shorter for high motor intent trials compared to low motor intent trials (F_(1, 43)_ = 48.00, p < 10^−8^, repeated measures ANOVA), and was unaffected by saccade latency (F_(2, 86)_ = 0.89, p = 0.41). In contrast, the neuronal saccade target selection latency in motor neurons was independent of the strength of motor intent (F_(1, 15)_ = 0.72, p = 0.41) but covaried with saccade latency (F_(2, 30)_ = 6.22, p < 0.01). Together, an increased level of motor intent toward a potential saccade location within the neurons’ RF enhances the spike responses of visuomotor neurons in the SC. This modulation occurs independently of the physical preparation or execution of the saccade. In contrast, SC motor neurons specifically represent the preparation or execution of the saccade itself, rather than the level of motor intent.

### Independent control of perceptual sensitivity and perceptual decision criteria

Previous studies have suggested that spiking of visuo-motor as well as visual neurons in SC closely varies with perceptual attentional performance and decision^15,19^. Thus, we examined how the activity of individual SC neurons relates to perceptual *d’* and decision criteria, independently in relation to response bias. Monkeys’ behavioral d’ and decision criteria were independently controlled using the same visual orientation change detection task as in Figure 1 (**Fig. 2a**; **Methods**).

**Figure 2.**
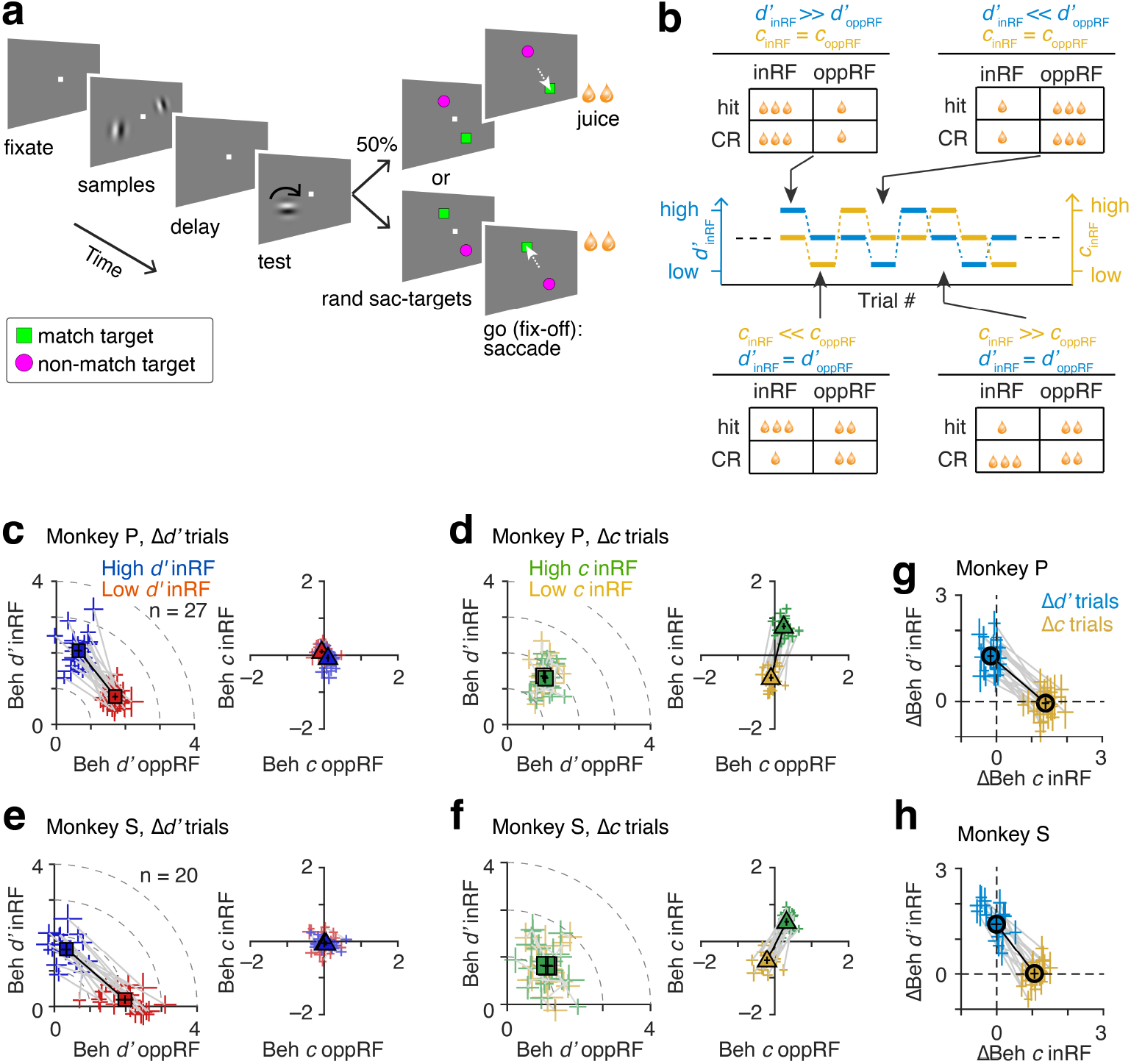
Independent control of behavioral sensitivity (*d’*) and perceptual criteria. (**a**) Visual orientation change detection task. The task was like that in Figure 1, exception the positions of the saccade targets were randomized on each trial. (**b**) Monkeys’ spatial attention and decision criteria were independently controlled between the two stimuli locations in interleaved blocks of trials by adjusting the reward size for correct detections. (**c, e**) Monkeys’ behavioral *d’* across sessions (monkey P, 27; monkey S, 20) for Δ*d’* trial blocks. Behavioral *d’* is better (*left*) on trials when the test stimulus appeared at the attended location (valid trials) compared to the unattended location (invalid trials) without any changes in criteria (*right*). Gray lines, individual session. Filled squares, session average. Error bars, 95% confidence intervals. (**d, f**) Similar to **Figs. 1c** and **1e**, except for Δ*c* trial blocks in same experimental sessions. Behavioral *d’* remained uniform (*left*), while criteria (*right*) changed between low and high values. (**g-h**) Changes in behavioral *d’* and criteria at one of the two locations contralateral to recorded SC (monkey P, right visual field; monkey S, left visual filed) across sessions on Δ*d’* and Δ*c* trial blocks. Gray lines, individual session. Error bars, 95% confidence intervals.

However, the locations of match and non-match saccade targets were randomly interchanged across trials. This dissociated animal’s spatially selective response bias or motor intent from perceptual *d’* and perceptual decision criteria as there was no prior information about the saccade targets until after the test stimulus was turned off. Animals were instructed to switch their spatial attention by adjusting reward sizes for correct detections between the two locations in small blocks of trials (monkey S, 120-160 trials, monkey P, 120-140 trials) (Δ*d’* trials, **Fig. 2b**). While a relatively large average reward for correct responses (hits and CRs) at one location motivated animals to selectively shift their behavioral *d’*, the reward size ratio for hits and CRs was varied somewhat across trials to maintain decision criteria close to zero. In separate blocks of trials, perceptual decision criteria was controlled between low and high values at a location independent of behavioral *d’* by adjusting the reward size for hits and CRs in small blocks of trials that were randomly interleaved with Δ*d’* blocks (Δ*c* trials, **Fig. 2b**).

Behavioral *d’* increased significantly at the attended location compared to the unattended location on Δ*d’* trial blocks (monkey P, 27 sessions F_(1, 52)_ = 17.38, p < 10^−9^; monkey S, 20 sessions, F_(1, 38)_ = 24.48, p < 10^−4^, repeated measures ANOVA; *left*, **Figs. 2c** and **e**), without significant changes in decision criteria (monkey S, F_(1, 38)_ = 0.95, p = 0.33, monkey P, F_(1, 52)_ = 1.44, p = 0.23; *right*, **Figs. 2c** and **e**). On Δ*c* trial blocks within the same experimental sessions, animals selectively shifted their decision criteria (monkey P, F_(1, 52)_ = 397.02, p < 10^−^18; monkey S, F_(1, 38)_ = 186.87, p < 10^−15^; *right*, **Figs. 2d** and **f**) without significant changes in behavioral *d’* (monkey P, F_(1, 52)_ = 1.47, p = 0.23; monkey S, F_(1, 38)_ = 0.05, p = 0.83; *left*, **Figs. 2d** and **f**). This independent control of perceptual *d’* and decision criteria (**Figs. 2g** - **h**) was crucial for investigating the relative contributions of distinct perceptual and behavioral components of attentional performance by SC neurons.

### Spike modulation of SC neurons associated with changes in perceptual sensitivity and decision criteria

We simultaneously recorded spiking activities from populations of visuo-motor (n = 165), visual (n = 88) and motor neurons (n = 52) in the SC while the monkeys’ behavioral *d’* and perceptual decision criteria were controlled during the visual attention task (monkey S, n = 127; monkey P, n = 178; **Supplementary Fig. S3**). Visuo-motor neurons strongly responded to the sample stimuli for the trial blocks when the monkey’s behavioral *d’* at the RF was high (*blue* and *orange*, **Figs. 3a-b**). However, spike rates did not change during changes in decision criteria of similar magnitude (*yellow* and *green*, **Figs. 3a-b**). There was a significant spike modulation as measured by neuronal *d’* during the sample stimuli presentation for behavioral Δ*d’*, but not for decision Δ*c* in visuo-motor neurons (for behavioral Δ*d’*, mean ± SEM = 1.29 ± 0.08; *P* < 10^−17^; for decision Δ*c*, mean ± SEM = 0.005 ± 0.06; *P* = 0.71; n = 105/165, signed rank test; **Fig. 3c**). A much weaker but significant neuronal modulation was seen in SC visual neurons for behavioral Δ*d’*, but not for behavioral Δ*c* (for behavioral Δ*d’*, mean ± SEM = 0.2 ± 0.06; *P* < 10^−02^; for decision Δ*c*, mean ± SEM = 0.023 ± 0.06; *P* = 0.4; n = 58/88, signed rank test; **Figs. 3d-e**). Spike rates of motor neurons were unaffected with behavioral Δ*d’* as well as decision Δ*c* (for behavioral Δ*d’*, mean ± SEM = 0.13 ± 0.21, *P* = 0.45; for decision Δ*c*, mean ± SEM = –0.19 ± 0.16, *P* = 0.14; n = 30/52, signed rank test; **Supplementary Fig. S4**).

**Figure 3.**
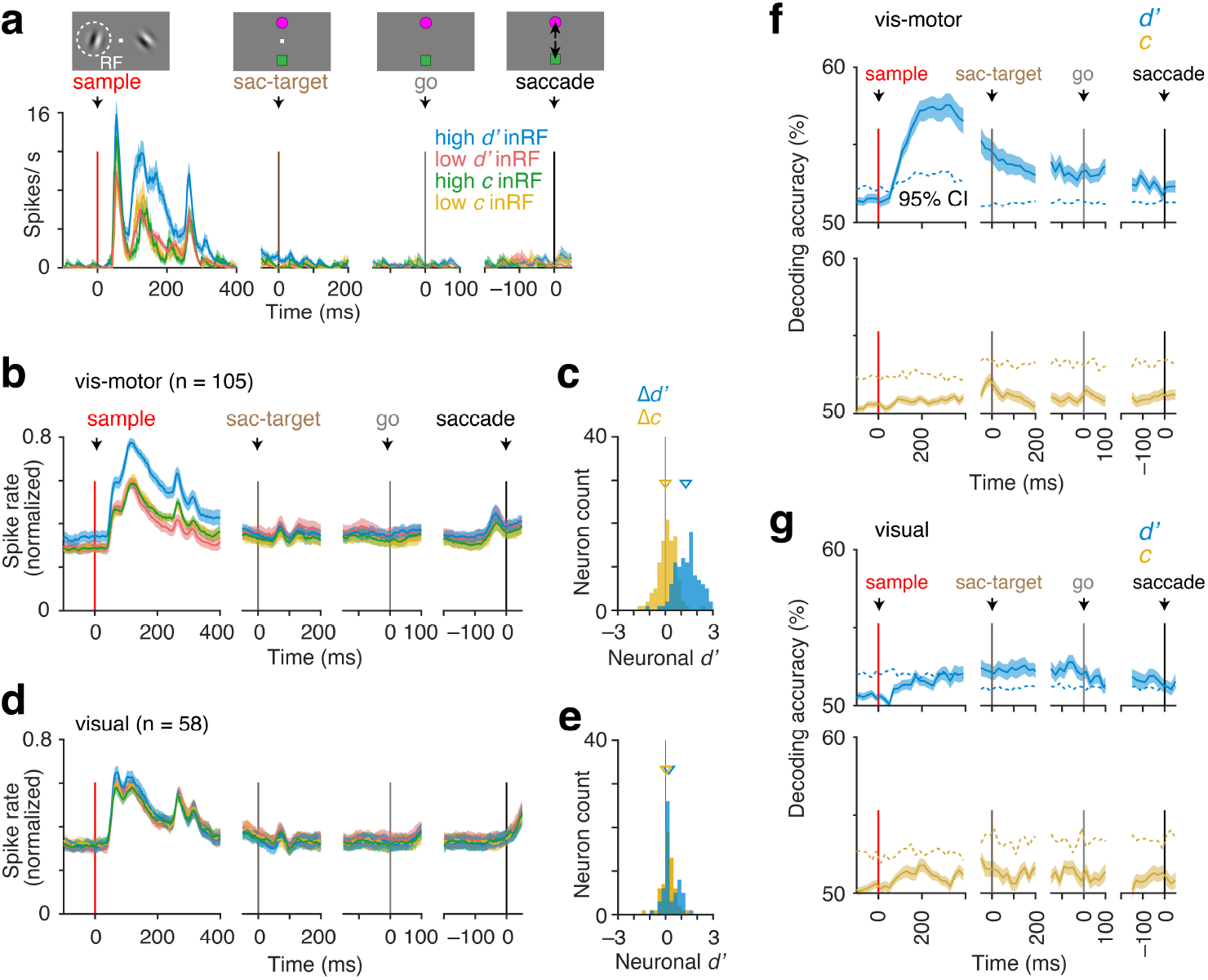
Neuronal modulation of SC neurons with changes in behavioral *d’* and perceptual decision criteria. (**a**) Trial averaged PSTHs of an example visuo-motor SC neuron when the monkey’s behavioral *d’* and decision criteria at the neuron’s RF location were independently controlled relative to the stimulus at the opposite hemifield (**Fig. 2**). The spike rates are aligned to the onsets of sample stimuli, saccade targets, go-cue and the saccade (*top*). Error bars, ±1 SEM. (**b**) PSTHs are similar to those in (a), except for the population of visuo-motor neurons when one of the visual sample stimuli was located within the neuron’s RF (n = 105/165). (**c**) Spike modulation associated with behavioral Δ*d’* and Δ*c*, measured by neuronal *d’*. Triangles represent population means. (**d-e**) Similar to (b-c), except for visual neurons in the SC (n = 58/88). (**f-g**) Decoding accuracies for behavioral *d’* (*top*) and decision criteria (*bottom*) from spike counts using a linear classifier based on support vector machine, averaged across individual visuo-motor (f) and visual (g) neurons, as shown in (b) and (d). Dashed lines, 95% confidence intervals based on shuffled trials. Error bars, ±1 SEM.

To further quantify how the activity of individual SC neurons relates to perceptual *d’* and decision criteria independently, we examined decoding accuracy (10-fold cross validation) using a linear classifier based on a support vector machine (**Figs. 3f-g**). Spike counts of visuo-motor neurons reliably encoded information about behavioral *d’*, but not decision criteria, during the sample stimuli, compared to visual and motor neurons, despite the equivalent magnitudes of the monkeys’ behavioral Δ*d’* and Δ*c* (**Figs. 2g-h, Supplementary Fig. S4**).

One possible reason for the absence of perceptual Δ*c*-related neuronal modulation in SC neurons during the sample stimuli could have been that the saccade targets were positioned far beyond the neuron’s response field. Because the criterion is closely related to selecting a response choice, criterion-related neuronal processing might have been associated with the choice targets rather than the attended sensory stimuli in trials when Δ*c* was manipulated. To test this possibility, we measured Δ*d’* and Δ*c* -related modulation of SC neurons whose response fields overlapped with one of the saccade targets while controlling monkeys’ *d*’ and *c* independently within the same experimental sessions, as shown in Figure 2 (**Supplementary Fig. S5**). There was no significant spike modulation for Δ*c* in any of the neuron groups during the sample stimulus (mean ± SEM neuronal *d’*, visuo-motor, 0.005 ± 0.07; *P* = 0.38; visual, 0.012 ± 0.14; *P* = 0.4; motor, –0.20 ± 0.17; *P* = 0.43; mean ± SEM neuronal mod. index, visuo-motor, -0.006 ± 0.01; *P* = 0.46; visual, 0.0 ± 0.01; *P* = 0.46; motor, 0.016 ± 0.03; *P* = 0.29; signed rank test). Moreover, behavioral Δ*c* did not affect the spike response at the onset of saccade targets (mean ± SEM neuronal *d’*, visuo-motor, –0.23 ± 0.15; *P* = 0.17; visual, –0.06 ± 0.12; *P* = 0.43; motor, 0.03 ± 0.27; *P* = 0.44; mean ± SEM neuronal mod. index, visuo-motor, 0.005 ± 0.01; *P* = 0.25; visual, 0.01 ± 0.01; *P* = 0.31; motor, 0.006 ± 0.05; *P* = 0.32; signed rank test). Spike counts of these neurons also did not represent information about decision criteria during the sample stimuli or saccade target presentations, as measured by decoding accuracy in cross validated trials (10-fold) (**Supplementary Fig. S5**). However, information about behavioral *d’* was represented by visuo-motor neurons during the presentation of saccade targets. Therefore, a strong spike modulation of SC visuo-motor neurons was specifically linked to a change in perceptual *d’*, rather than perceptual decision criteria.

### Neuronal modulation in SC visuo-motor neurons varies with behavioral sensitivity and spatial alignment with attended location

Our analysis showed that the increase in spike rate of SC visuo-motor neurons with higher behavioral *d’* reflects a neuronal correlate of trial-averaged behavioral performance as spatial attention shifts between stimulus locations across blocks of trials. However, attentional states can fluctuate on much shorter timescales. Therefore, we next examined at a granular level how neuronal modulation covaries with both the graded levels of behavioral *d’* within individual sessions, and the spatial alignment between a neuron’s RF and the attended location.

We measured the proximity of SC visuo-motor neurons’ RFs to the attended Gabor sample stimulus using Mahalanobis distance in the experimental sessions described in Figure 2 (n = 129/165, **Fig. 4a-b**). The dependence of neuronal modulation on the spatial distance between the neuron’s RF and the attended stimulus, as well as on behavioral *d’* at the single-neuron level, was estimated using a linear regression model (**Fig. 4c)**. Neurons with RFs closer to the attended stimulus showed greater modulation (β1 = –0.1, p < 10^−3^; *left*, **Fig. 4c**). Additionally, neuronal *d’* was positively correlated with behavioral *d’* (β2 = 0.45, p < 10^−3^; *right*, **Fig. 4c**). At the population level, neuron’s spatial proximity to the attended location explained a greater proportion of variance in neuronal *d’* compared to behavioral *d’* (13.5% versus 7.4%; **Fig. 4d**).

**Figure 4.**
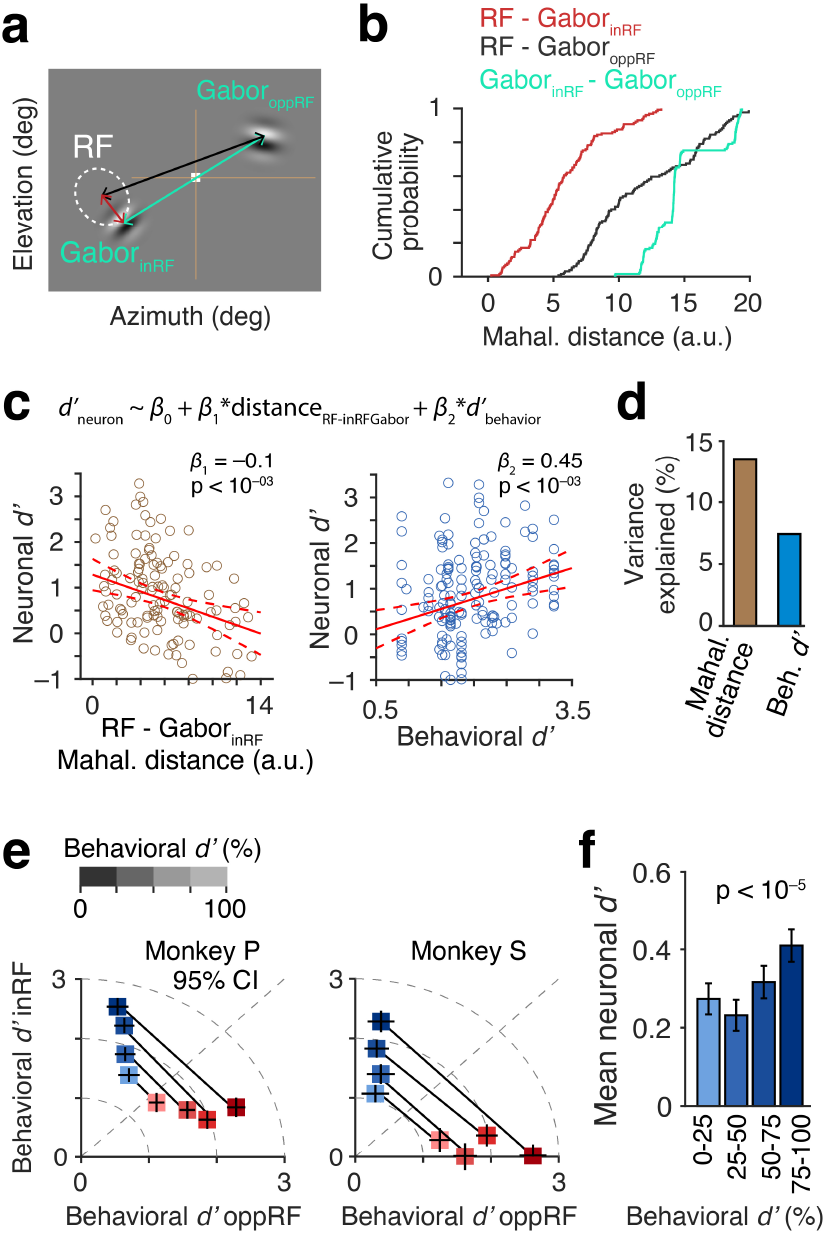
Neuronal modulation of SC visuo-motor neurons related to spatial attention depends on the level of behavioral sensitivity and the alignment of the neuron’s spatial response field with the attended stimulus location. (**a**) A schematic illustrating the alignment of a neuron’s spatial RF and Gabor sample stimuli as measured by Mahalanobis distance. (**b**) Cumulative distributions of Mahalanobis distances between visuo-motor neurons’ RF (neurons, n = 129) and Gabor stimuli in two opposite hemifields, and between the two Gabor stimuli (sessions, N= 47). (**c**) Linear regression fit for neuronal d’ as a function of the average behavioral *d’* (*Right*) and the Mahalanobis distance between the RF and the Gabor stimulus in the same hemifield (*Left*, RF-Gabor_inRF_). (**d**) Variances explained by the RF-Gabor_inRF_ distance (MD) and behavioral d’ in the regression fit in (c). (**e**) Trials within each session were sorted according to behavioral d’ into equal quantiles, and then plotted for the two monkeys. (**f**) Trial-averaged neuronal d’s as a function of quantiled trial groups with varying behavioral d’.

To assess behavioral performance on a shorter timescale, behavioral *d’* was computed using a 25-trial moving window separately for all four trial types (low versus high selective attention; in-RF versus opposite-RF location) during Δ*d’* manipulation trial blocks within each session. These behavioral *d’* distributions were sorted into four equal quantiles to get a within-session gradient of behavioral performance (**Fig. 4e**). Neuronal *d’* scaled with behavioral *d’*, with stronger neuronal modulation observed in high-performing trials (F_(3, 492)_ = 9.04, p < 10^−5^, repeated measures ANOVA; **Fig. 4f**). These results suggest that SC visuo-motor neurons integrate spatial and behavioral factors, with attentional modulation being stronger when neuron’s RF aligns with behaviorally relevant stimulus and attention is more intense.

### Single-trial decoding of distinct cognitive and behavioral factors associated with attentional performance from SC visuo-motor neurons

Previous studies, along with our results, suggest that different sensory, cognitive and behavioral factors can be multiplexed in single-trial spike trains of SC visuo-motor neurons. Thus, we decomposed single-trial population firing rates of SC visuo-motor neurons into demixed principal components to quantify relative encoding of motor response criteria (low versus high intent to select a saccade location; **Fig. 1**), perceptual decision criteria (low versus high bias to select saccade target independent of location; **Fig. 2**), perceptual *d’* (in-RF versus opposite-RF; Fig. 2), saccade location (in-RF versus opposite-RF; **Figs. 1** and **2**) and correct detection (correct versus error; Figs. 1 and 2) (**Fig. 5**; **Methods**).

**Figure 5.**
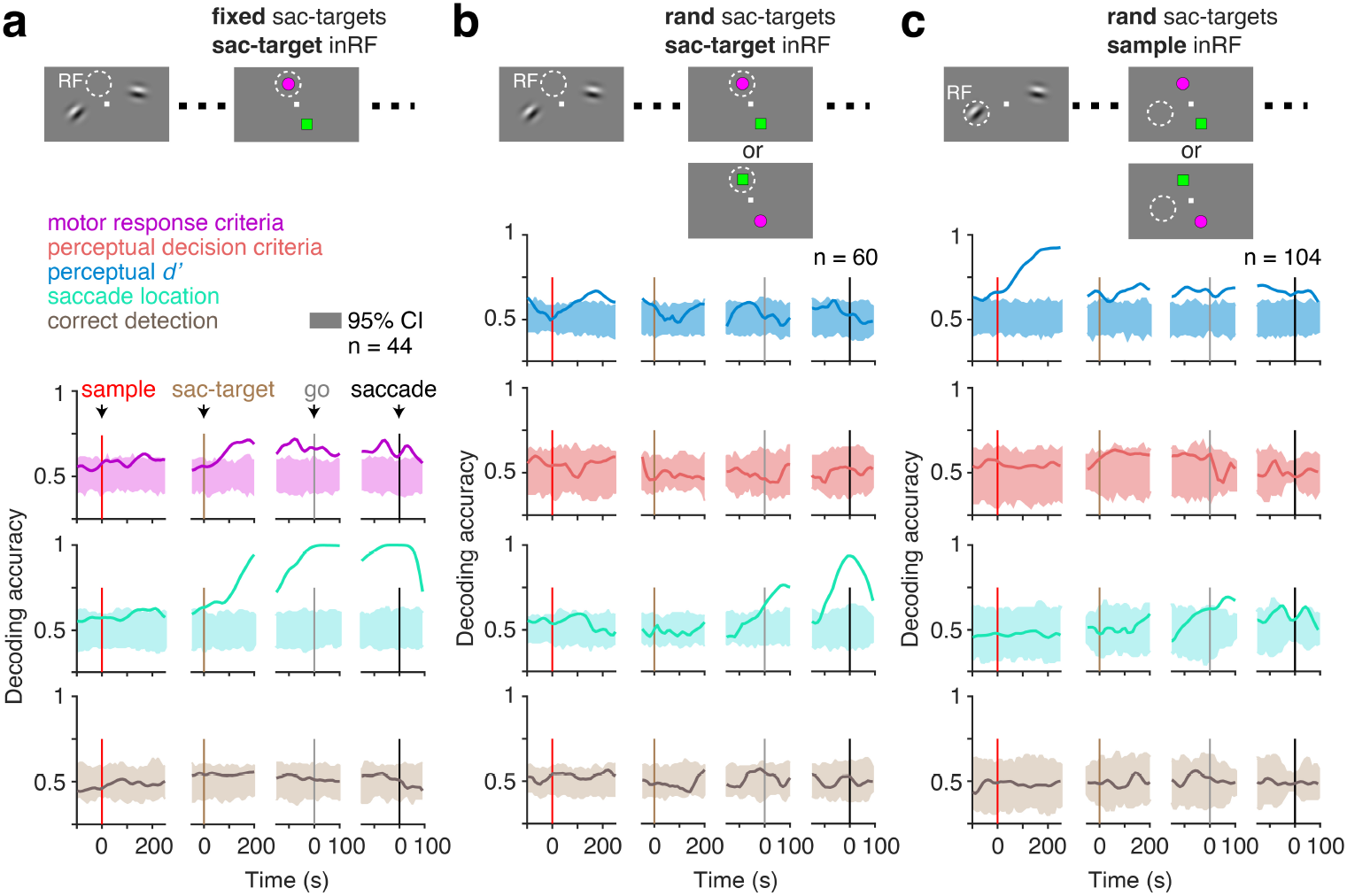
Single-trial decoding of distinct perceptual, cognitive and behavioral factors associated with attentional performance from SC visuo-motor neurons. (**a**) Time course of single-trial decoding accuracy (cross-validated leave-one out trials) for motor response criteria (response bias), executed saccade location and correct detection based on demixed principal components of population PSTHs of SC neurons as in Figure 1 (**Methods**). The locations of the saccade targets were fixed, with one target placed within the neuron’s RF. Only neurons for which at least 10 trials were conducted for each decoding factor were included. Error bars, 95% confidence intervals from shuffled trials. (**b**) Similar to (a), except for sessions in Figure 2 where one of the saccade targets was placed within the neuron’s RF and the saccade targets were randomized (n = 60/165). Additional decoding factors included perceptual *d’* and perceptual decision criteria. (**c**) Similar to (b), except for the sessions when one of the sample stimuli was placed within the neuron’s RF (n = 104/165).

When saccade targets were within the neuron’s RF, decoding accuracy for saccade location increased sharply before saccade execution, and (*cyan traces*, **Figs. 5a-b**) and emerged earlier when saccade target locations were fixed (*cyan traces*, **Fig. 5a**). Perceptual *d’* was also strongly encoded during the sample period when the attended stimulus, rather than the saccade target, was presented at the neuron’s RF (*blue traces*, **Figs. 4b-c**). Information regarding the motor response criteria was represented independently of saccade execution when the saccade target locations were fixed (*magenta traces*, **Fig. 4a**). In contrast, perceptual decision criteria (*red traces*, **Figs. 4b-c**) as well as correct detection (*brown traces*, **Figs. 4b-c**) were not significantly encoded during the trial. These results suggest that SC visuo-motor neurons multiplex perceptual, response selection and motor execution depending on instantaneous task demands.

## DISCUSSION

Our findings demonstrate that neuronal activity in the superior colliculus (SC) is strongly modulated by perceptual sensitivity (*d’*) associated with visual spatial attention. The magnitude of this modulation depends on the spatial proximity of the locus of attention to the neuron’s response field (RF), as well as the intensity of perceptual *d’*. However, SC activity does not reflect whether the monkeys correctly detected a stimulus change at the single-trial level. Instead, the neuronal modulation corresponds to an elevated cognitive state associated with improved perceptual *d’*. The SC activity was also enhanced with lower motor response criteria, which occurred when there was a bias to select a saccade target located within the neuron’s RF. In contrast, shifts in perceptual decision criteria at the neuron’s RF, which involved a bias toward selecting a target category rather than location, did not affect SC neuronal activity.

Lue and Maunsell ^9,20^ showed that neurons in the lateral prefrontal cortex (LPFC) represent both behavioral *d’* and criterion, while the visual area V4 selectively represents behavioral *d’* in a similar visual detection task. However, the motor component was not dissociated from the perceptual decision. By behaviorally dissociating these components, we reveal a disconnect between SC activity and perceptual performance. First, SC neuronal activity was independent of perceptual decision, challenging the proposed role of SC in perceptual decision making. Second, single-trial perceptual detection was independent of SC activity, instead representing a cognitive state of enhanced spatially selective perceptual sensitivity, as observed in visual cortex^8,9^.

### Does the oculomotor activity in the SC support a common cognitive function of ‘selection’?

Results from various experimental tasks investigating the functions of the SC suggest that its neuronal activity mediates many different cognitive functions, including spatial orientation, target selection, covert attention, and saccadic eye movements. Consistent with our findings, previous research shows that SC visuomotor neurons respond to visual target selection in their response field (RF) independently of saccade execution^17^. Deficits in spatial target selection, as observed in visual search tasks ^21^, may underlie ipsilateral choice biases or delayed decisions when the SC is unilaterally inactivated in two-alternative forced-choice tasks ^11,22^.

SC activity is also crucial for behavioral performance in covert visual attention tasks that rely on perceptual sensitivity with inactivation–either pharmacological or optogenetic, impairing perceptual performance^10,13,23^. In light of these previous findings, our results on perceptual Δ*d*’- and response Δ*c*-related SC modulation support the role of SC in a common ‘spatial selection’ mechanism that either enhances the perception of stimulus details or guides action accuracy (or response time) in a task-dependent manner. Selective attention is often broadly defined as the common mechanism for selective prioritization ^2,24^ of a stimulus for perceptual benefit, working memory, or motor action. However, because perceptual decision Δ*c* did not modulate SC activity, our findings suggest that the SC’s oculomotor circuitry does not support the ‘selection’ of non-spatial content, highlighting a fundamental distinction between spatial and non-spatial decision processes in the SC.

### SC interaction with other brain structures in mediating functions beyond oculomotor control

Recent studies have shown that the visual neurons in the SC represent higher-order cognitive functions, specifically object category ^25^. This suggests that specific microcircuitry of SC neuronal subpopulations can support distinct neuronal computations beyond the oculomotor controls depending on the task-demands ^16,23,26^. Brain areas reciprocally connected to the SC, such as the sensory visual cortex, intraparietal cortex (LIP), and frontal eye field (FEF), are known to represent different aspects of cognition and decision-making ^27-29^. Independent controls of these cognitive factors, combined with specific manipulations of the activity of neurons or circuits within these areas, may provide insights into the causal contributions of the SC to cognition and behavior.

### Information readout by brain structures downstream to the SC, whether to perceive in detail or to act upon

SC is a node among within a network of many brain structures that mediate distributed computations related to visual attention, decision formation and motor action. The question arises regarding how the neurons downstream of the SC decode an enhanced spiking if the same neuron responds to perceptual sensitivity, response selection and motor planning. Previous studies show that downstream neurons in the SC-FEF pathway via mediodorsal thalamus differentially process complex sensory and motor signals, with thalamus filtering out delay activity and supporting FEF in predominantly retaining perisaccadic activity ^30^. Simultaneously recorded large ensembles of SC neurons reveal that population coding structures sensory and motor-related activities in distinct patterns, enabling downstream areas to decode them, despite individual neurons responding to multiple cognitive and motor processes ^31^. Pairwise correlation, an information encoding strategy independent of spike rates, is sensitive to distinct neural processes guiding saccades and varies within and between SC neuron subclasses ^26^. This may also contribute to distinguishing multiplexed signals by regulating information transfer based on task context.

## Conclusion

The SC strongly represents a cognitive state associated with improved perceptual sensitivity in visual spatial attention tasks, independent of response bias. However, signals related to non-spatial perceptual decisions and trial-by-trial perceptual detection are not represented in the SC when spatial-component is dissociated.

## METHODS

### Animals and surgery

Two adult male rhesus monkeys (*Macaca mulatta*, 9 and 13 kg) were each implanted with a titanium head post using aseptic surgical techniques before behavioral training began. After the completion of training (3 to 5 months), we surgically implanted a stainless-steel recording chamber targeting the SC on one side (*right*, monkey S; *left*, monkey P) to access the SC, guided by MRIs obtained before the initial surgery. The cylinders were centered on the skull at 3.5 mm A, 10.0 mm L, and tilted in the coronal plane to advance toward the midline (monkey P, 9°; monkey S, 11°). The same two monkeys were used in previous studies that described different findings on neuronal responses in area V4 ^8,32^ and locus coeruleus ^18^. All experimental procedures were approved by the Institutional Animal Care and Use Committee of the University of Chicago and followed the U.S. National Institutes of Health guidelines.

### Visual change-detection task

Monkeys sat in a primate chair facing a calibrated CRT display (1024 × 768 pixels, 100 Hz frame rate) at 57 cm viewing distance inside a darkened room. Binocular eye position and pupil area were recorded at 500 Hz using an infrared camera (EyeLink 1000, SR Research). Trials started once the animal fixated within 1.5° of a central white spot (0.2° square) presented on a mid-level gray background (**Fig. 1a**) ^18^. The animal had to maintain fixation until its saccade response at the end of the trial. After a randomly selected fixation period of 400 to 700 ms, two achromatic Gabor sample stimuli appeared for 200 ms, one in each visual hemifield. After a random variable delay of 200 to 300 ms, a Gabor test stimulus appeared for 200 ms at one of the two stimulus locations, selected randomly with equal probability. Shortly after the test stimulus turned off (100 to 150 ms), two saccade targets of different color and shape (non-match target, green square; match target, magenta circle; 0.30°-0.4°) appeared in opposite directions along an imaginary line orthogonal to the axis of the samples. A go-signal (fixation spot turning off) occurred 150 to 200 ms after the saccade target appeared and indicated the animal should make a saccade to the appropriate saccade target depending on the change in the test Gabor orientation relative to the sample. The orientation of the test stimulus changed from the sample stimulus on random half of trials (non-match trial; 32° for monkey S and 42 ° monkey P). On the rest half of the trials, the orientation of the test stimulus remained unchanged (match trial). Orientation of sample Gabor stimulus at each location was randomized across trials from 0° to 175° (5° intervals). Orientations of left and right sample stimuli were independent and never identical. Gabor stimuli were varied each day and remained unchanged throughout each session (monkey P, left Gabor, azimuth, −9.0° to −1.0° [median = −6°, inter-quartile = −7.0°, −3.7°], elevation, −8.0° to 7.0° [median = −0.85°, inter-quartile = −4.9°, 4.7°]; sigma, 0.75° to 2.7° [median = 1.85°, inter-quartile = 1.1°, 2.4°], spatial frequency, 0.15 to 0.5 [median = 0.3, inter-quartile = 0.18, 0.4] cycles per degree, right Gabor, azimuth, 1° to 13° [median = 7°, inter-quartile = 5°, 11.6°], elevation, −9.1° to 9.6° [median = −4°, inter-quartile = −7.7°, 7°], sigma, 0.65° to 3.0° [median = 2.4°, inter-quartile = 1.3°, 2.7°], spatial frequency, 0.14 to 2.2 [median = 0.27, inter-quartile = 0.17, 0.4] cycles per degree; monkey S, left Gabor, azimuth, −15° to −2.3° [median = −7°, inter-quartile = −10°, −6.5°], elevation, −10° to 7.5° [median = –1°, inter-quartile = −1°, 3°], sigma, 1° to 1.8° [median = 1.2°, inter-quartile = 1.15°, 1.5°], spatial frequency, 0.3 cycles per degree, right Gabor, azimuth, 2.3° to 8.0° [median = 7°, inter-quartile = 6°, 7°], elevation, −7.5° to 7° [median = 1°, inter-quartile = −1.8°, 1°], sigma, 1° to 1.3° [median = 1.2°, inter-quartile = 1.15°, 1.2°], spatial frequency, 0.3 cycles per degree). Targets’ eccentricities remained unchanged throughout each session. (monkey P, 3.6° to 5.7° [median = 4.3°, inter-quartile = 4.1°, 4.6°]; monkey S, 3.3° to 9° [median = 4.1°, inter-quartile = 3.6°, 4.3°]).

### Independent control of behavioral *d’* and criterion associated with visual spatial attention

Monkeys’ behavioral response criterion (*c*) (**Figs. 1a-b**) in one of the sample stimulus locations was controlled between low and high values across interleaved trial blocks, while behavioral sensitivity (*d’*) between the two stimulus locations remained uniform by adjusting reward size for correct detections. The locations of match and non-match saccade targets remained fixed throughout a trial block (120-160 trials), and were cued by a few instruction trials (10-15) that preceded the trial block start and were not included in the analysis. The probability of test trials for the two sides was equal. The ratio of the reward size for hit trials over correct-rejection trials (hit:CR) associated with one of the sample stimuli was 2-3 to achieve a low response criterion (higher bias to select the non-match saccade target). Conversely, a hit:CR reward ratio of 0.3-0.5 led to a high response criterion (lower bias to select the non-match saccade target). Both the reward ratio for hit:CR in the opposite sample stimulus location, as well as the average reward size (hit and CR) between the two stimulus locations, were kept near 1 throughout the session. This maintained a uniform criterion at the opposite location and uniform behavioral d’ between the two locations (**Figs. 1b**).

Perceptual decision criterion were independently controlled between low and high values (four trial conditions) in interleaved blocks of trials (120-160 trials, **Fig. 2a**). The criterion was controlled in the same manner as described in the previous section for controlling the response criterion, except that the match and non-match saccade targets randomly switched their positions across trials. This dissociated the spatially selective response preference, such that the criterion was a perceptual decision criterion that was independent of spatial ‘selection’ for saccade or selective perceptual *d’*.

Behavioral *d’* was controlled between the two locations using both different reward size as well as different probability of test trials for the two sides ^18^ (**Fig. 2a**). Average reward size ratio (hit and CR) for the valid location (attended) over the invalid location was 3.0-3.5. The ratio of valid and invalid test stimulus probabilities for the locations was 2-3. Both the test stimulus probability and reward size contributed to reliably control the behavioral *d’* at the two stimulus locations. The hit:CR reward ratio at the two locations was 1. Varying reward size was particularly effective in keeping the perceptual decision criterion close to zero. In previous studies, we have used reward size differences with and without a test stimulus probability difference, to control behavioral *d’* and criterion values ^8,9^. Expecting a high reward size or a high expectation of a test stimulus might access a common valency, which is the product of the probability of a test stimulus and its reward size, associated with a stimulus location that drives the motivation to shift attention ^33^.

### Memory guided saccade task

In each session, we mapped visual and saccade response fields by having the monkeys do a memory-guided saccade task before the visual attention task (**Fig 1f**). After fixating for 400 to 600 ms, a circular saccade target (0.35° diameter) appeared for 250 ms at one of the six equally spaced peripheral locations. A common target eccentricity was selected based on recorded neurons’ visual response fields (2.5°-10°). Following a delay of 500-800 ms, the fixation spot disappeared, and the monkey executed a saccade to the remembered location of the visual target (within 4°-7°). At least 20 correct trials were collected at each stimulus location in each session.

### Neurophysiological recordings

Neuronal signals from an extracellular multielectrode linear array (16 channel V-probe and 32 channel U-probe; Plexon Inc.) were amplified, bandpass filtered (250 to 7,500 Hz), and sampled at 30 kHz using a data acquisition system (Cerebus, Blackrock Microsystems) (**Figs. 1** and **3**). We simultaneously recorded from multiple single units and small multiunits over 56 sessions (25 sessions for monkey S; 31 sessions for monkey P).

### Data analysis

#### Behavioral performance

All completed trials were included in our analysis. Behavioral *d’* and perceptual criterion (*c*) at a spatial location were measured from hit rates within nonmatch trials and false alarm (FA) rates within match trials using 1-dimensional signal detection theory^7,32^ as: 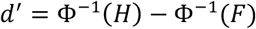 and 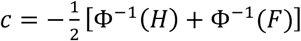 where Φ–1 is inverse normal cumulative distribution function; H and F are respectively the rates of hits and false alarms.

#### Neuronal response

Spikes from each electrode were sorted offline (Offline Sorter, Plexon Inc.) by manually well-defining cluster boundaries using principal components analysis and waveform features. Well-isolated clusters were classified as single units from multiunits based on the isolation quality of unit clusters. The degree to which unit clusters were separated in two-dimensional (2D) spaces of waveform features (first three principal components: peak, valley, and energy) was measured by multivariate analysis of variance (MANOVA) *F* statistic using Plexon Offline Sorter (Plexon Inc.). A unit cluster of MANOVA *P* < 0.05 was considered as a single unit that indicates that the unit cluster has a statistically different location in 2D space and that the cluster is statistically well separated. Spike counts in 2 ms bins were smoothed using half-Gaussian kernel (standard deviation 15 ms, rightward tail). Spike trains were aligned to the onsets of the stimulus (visual stimulus and saccade target) or task events (fixation, go cue, saccade) across trials to generate peristimulus time histograms (PSTHs) for spike rates. Population spike rates were calculated by averaging individual unit PSTHs after normalizing them to their peak spike rate.

#### Classification of SC neuronal responses

Neurons were categorized based on their average evoked spike counts to visual targets (50 to 300 ms after onset) and saccades (−100 to 50 ms after the saccade’s onset), during the memory-guided saccade task (**Fig. 1f, Supplementary Figs. S1** and **S3**). An evoked spike rate was considered significant if it was greater than (p < 0.05, signed rank sum test) the response during the fixation period (−250 to 0 ms from the onset of the visual target). Neurons classified as visually selective (‘visual’) were identified based on a significant difference in their peak visual response (preferred response field) from the response to the visual target at the opposite location (180°, non-preferred response field; p < 0.05, signed rank sum test), and they had no saccade response. Similarly, neurons were classified as saccade selective (‘motor’) if their peak saccade-related response (preferred response field) was greater compared to the saccade response to the opposite location (180°, non-preferred response field; p < 0.05, signed rank sum test), and they exhibited no visual response. Neurons exhibiting both visual and saccade selectivity were categorized as “visuo-motor” neurons.

#### Neuronal d’

Neuronal spike rate modulations associated with behavioral Δ*d’* and Δ*c* were measured using neuronal sensitivity as: 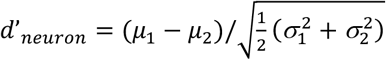 where *µ*_*i*_ and *σ*_*i*_ are the average and standard deviation of spike counts within 50 to 250 ms from the onset of sample stimuli or saccade targets (*i*, trial conditions, for Δ*d’* and Δ*c*, high or low) (**Figs. 3c** and **3e, Supplementary Figs. S4 and S5**). PSTHs for neuronal *d’* were calculated from spike count trains that had been smoothed using a Gaussian window with a sigma of 50 ms, binned at 20 ms intervals (10 ms overlap) (**Figs. 1i-j**).

#### Neuronal latency in saccade target discrimination

The earliest time at which neuronal response could reliably discriminate the location of monkey’s saccade (in-RF versus opposite-RF) was the neuronal latency for saccade target discrimination of a single unit (visuo-motor or motor; **Figs. 1k-l**). All trials within each session of the visual task described in Figure 1 were grouped into four quartiles based on the saccade latency relative to the go-cue. These quartiles were separately determined for saccades directed towards the in-RF or opposite-RF location. Single trial spike counts were smoothed using a Gaussian window with a sigma of 50 ms, binned at 20 ms intervals (10 ms overlap), and aligned relative to the go-cue. Subsequently, the area under the receiver operating characteristic (AUROC) curve was estimated for saccades directed towards the in-RF versus the opposite-RF for each neuron and each set of trial quartiles. The earliest time relative to the go-cue when the AUROC was significantly higher compared to the AUROC derived from shuffled trials (95% confidence interval, 100 repetitions) was determined as the neuronal latency for discriminating a saccade target to be in-RF versus opposite-RF.

#### Linear decoder for behavioral Δd’ and Δc

We used linear classifiers based on support vector machine to quantify single-trial encoding of behavioral Δ*d’* and Δ*c* in SC neurons (**Figs. 3f-g, Supplementary Figs. S4 and S5**). Single trial spike counts were binned at 50 ms intervals (30 ms overlap) and aligned to visual stimuli or task-related events. These counts were then sorted into two groups based on low or high values of behavioral d’ or criterion at the neuron’s RF. Classifiers were trained on the training dataset (leave-one-out, 10-fold, 1000 repetitions) and decoding accuracy to behavioral d’ or criterion on single trials was estimated on the cross-validation trials. Similar decoding accuracy was estimated on shuffled trials to obtain 95% confidence intervals (100 repetitions).

#### Proximity of neuron’s response field (RF) to attended sample stimulus

Proximity between each neuron’s RF and sample stimulus (**Fig. 4a-b**) was estimated by a modified Mahalanobis distance ^34^. It measures the statistical distance between two multivariate normal distributions similar to the Mahalanobis distance, except that it takes into account the covariance of both distributions: 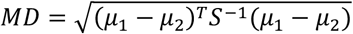 where S = (S_1_ + S_2_)/2; *µ*_i_ and S_i_ are respectively the mean and covariance of distribution *i* (1 and 2); ‘*T*’ and ^‘*–1*’^ are transpose and inverse respectively. For each neuron, the spatial RF was measured and fit using a bivariate Gaussian. This measure of the RF-stimulus proximity accounts for both the alignment of the stimulus with the neuron’s RF and the correspondence between stimulus and RF size.

#### Linear regression: neuronal d’–behavioral d’–RF to Gabor distance

A linear regression model (**Fig. 4c**) was used to fit neuronal 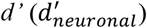 during sample stimuli (50-250 ms from the onset) to the behavioral *d’* at the neuron’s RF 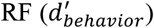 and the proximity between the neuron’s RF and the attended stimulus 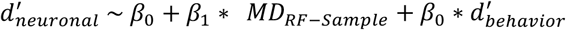 where β_i_ is regression coefficient (i = 0, 1 and 2).

#### Within-session correlation: spike count–neuronal modulation

For the dataset presented in Figure 2, behavioral *d’* was calculated over a sliding window of 25 trials (with a 5-trial overlap) for each of the four trial types (low versus high behavioral *d’*; in-RF versus opposite-RF stimulus location, total eight trial types) during the behavioral Δ*d’* manipulation trial blocks within each session (**Figs. 4e-f**). These behavioral d’ values for each trial type were then categorized into four equally spaced quartiles in each experimental session (**Fig. 4e**). Neuronal *d’* was calculated based on the spike counts during the sample stimuli (50-250 ms from the sample onset) for each of the four behavioral Δ*d’* quartiles (**Fig. 4f**).

#### Population decoding using demixed principal components analysis

Population activity of visuo-motor neurons in the SC was decomposed into distinct components, each carrying information about a single task-related or cognitive variable, using demixed principal component decompositions ^8,35^ (**Fig. 5**). Mean-subtracted and trial-averaged spike trains of each neuron were decomposed into marginalized averages, corresponding to task variables and a noise term. This ensures uncorrelated components. The least-square method minimizes a loss function penalizing the difference between marginalized and reconstructed data. Reconstructed data are projected onto a low-dimensional latent space and reconstructed with encoders. For the dataset in Figure 1, trials were categorized by motor response criteria (low versus high saccade bias in-RF), saccade direction (in-RF versus opposite-RF) and stimulus detection (correct [hit, CR] versus error [miss, FA]), resulting in six trial configurations (**Fig. 5a**). For the dataset in Figure 2, trials were classified by behavioral *d’* (low versus high in-RF), perceptual decision criteria (low versus high saccade bias), saccade direction, and stimulus detection, yielding eight configurations (**Fig. 5b-c**). All SC visuo-motor units with at least 10 trials for each condition from both monkeys were included in this analysis. Single-trial spike rates were filtered using a half Gaussian kernel (sigma = 30 ms) and then subsampled at 100 Hz. We analyzed spike rates over 500 ms (50 time points, starting 100 ms before sample onset, 50 ms before saccade target onset, 200 ms before go-cue, 400 ms before saccade onset). Decomposition into demixed components was performed on the training datasets using a leave-one-out approach and 200 repetitions. Subsequently, decoding accuracy was calculated on the remaining cross-validated test trials using the top three components. Confidence intervals (95%) were calculated on shuffled datasets.

#### Statistical analysis

Unless otherwise specified multifactor repeated measures ANOVA for comparing normally distributed datasets. Normality was checked using a Kruskal-Wallis test.

### Data availability

All data are available in the main text or the supplementary information. All behavioral and neuronal data analysis was done using Matlab (MathWorks Inc.). Behavioral task was controlled using custom-written software (https://github.com/MaunsellLab/Lablib-Public-05-July-2016.git).

