## Supplemental Figures for "Neuronal modulation of the superior colliculus associated with visual spatial attention represents perceptual sensitivity, independent of perceptual decision and motor biases"

### Supplementary Figures

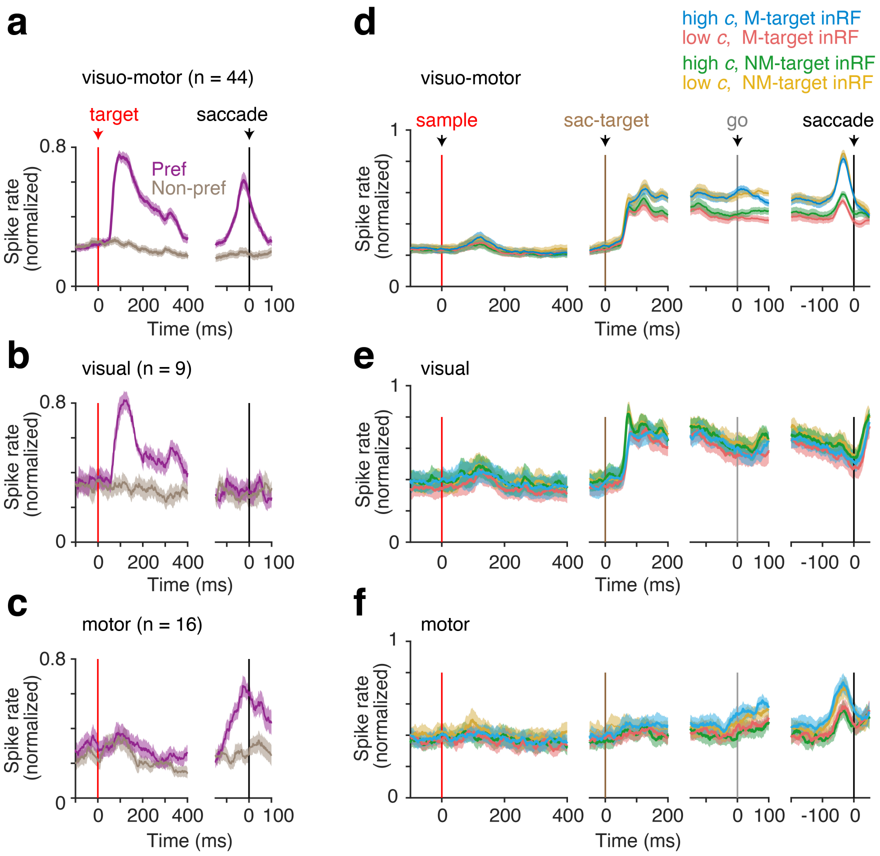


**Figure S1. Spike modulation of SC neurons associated with changes in response criteria where one of the saccade targets was within the neuron's response field (RF), and target positions remained fixed as in Figure 1. (a-c)** Neurons were classified into visuo-motor, visual, and motor neurons for the dataset in Figure 1. using delayed memory saccade task. Population-averaged PSTHs of visuo-motor (a), visual (b) and motor neurons (c) in monkey S and monkey P are aligned to visual target onset and saccade onset at the preferred and non-preferred RFs. Error bars represent ±1 SEM. **(d-f)** Population PSTHs of visuo-motor (d), visual (e) and motor-neurons (f) as in (a-c) during visual orientation change detection task in Figure 1. Spike rates are aligned to onsets of sample stimuli, saccade targets, go cue (fix-off) and saccade. Error bars, ± 1 SEM.

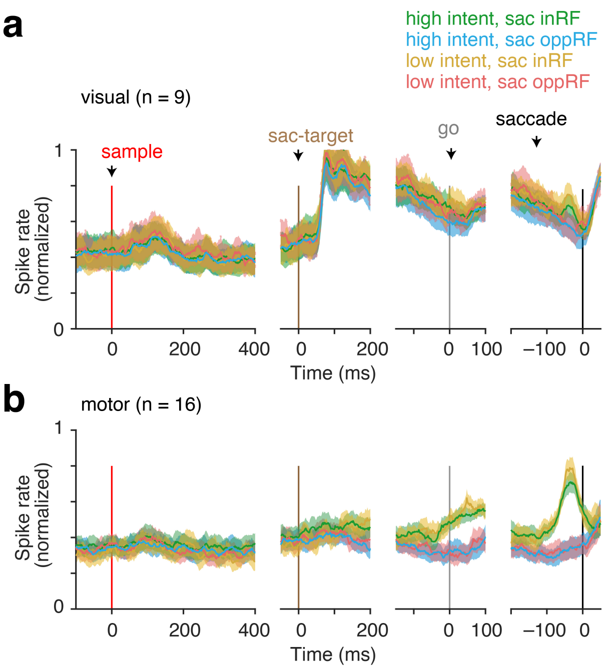


**Figure S2. Spike modulation of SC neurons associated with changes in response bias (motor intent), when one of the saccade targets was placed within the neuron's response field (RF), and target positions remained fixed as in Figure 1. (a-b)** Population PSTHs of visual (a) and motor neurons (b), grouped by saccade location and response bias level (saccade target selection) during visual orientation change detection task in Figure 1. Spike rates are aligned to onsets of sample stimuli, saccade targets, go cue (fix-off) and saccade. Error bars, ± 1 SEM.

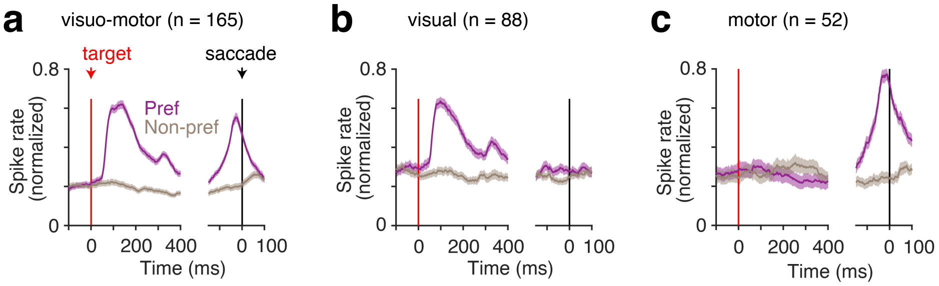


**Figure S3. Classification of SC neuronal responses using memory-guided saccade task for the data set shown in Figure 3. (a-c)** PSTHs of visuo-motor (a), visual (b) and motor (c) neurons in the SC, aligned to the onset of visual target and saccade for neuron's preferred and non-preferred RFs. Error bars, ± 1 SEM.

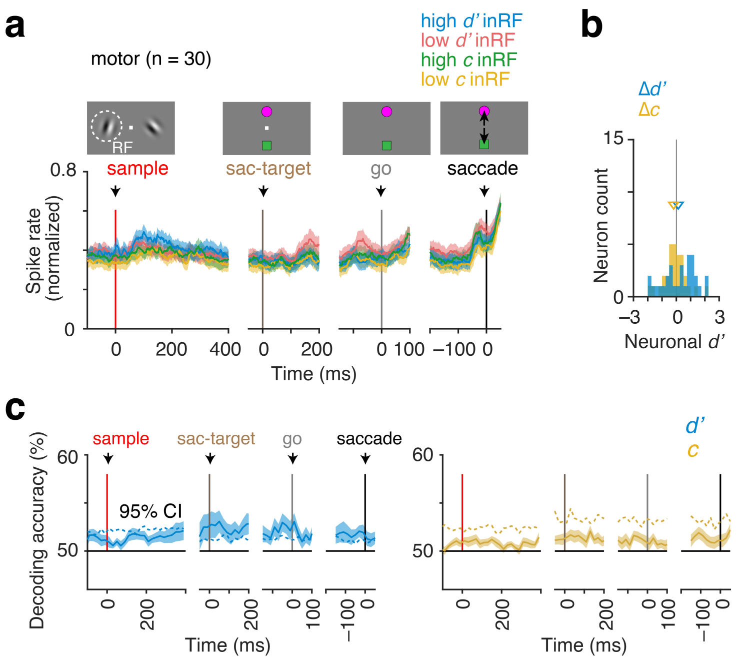


**Figure S4. Neuronal modulation in SC motor neurons with behavioral $\Delta d'$ and perceptual decision $\Delta c$, when the one of the sample stimuli (randomized) was placed in the neuron's RF.** (**a-b**) Population PSTHs of SC motor neurons when monkeys' spatial attention and decision criteria were independently controlled, and one of the sample stimuli was placed within the neuron's RF (n = 30/52). The spike rates are aligned to the onsets of sample stimuli, saccade targets, go-cue and the saccade. Error bars, ± 1 SEM. (**b**) Neuronal modulation as measured by neuronal $d'$ based on spike-counts over 200 ms (50-250 ms from the onset of sample stimuli, associated with changes in behavioral $d'$ ($\Delta d'$) and decision criteria ($\Delta c$). (**c-d**) Similar to (a) and (b), for visual neurons. (**c**) Decoding accuracies for $\Delta d'$ (*left*) and $\Delta c$ (*right*) based on SVM linear classifiers, averaged across individual motor neurons. Dashed lines, 95% confidence intervals based on shuffled trials. Error bars, ± 1 SEM.

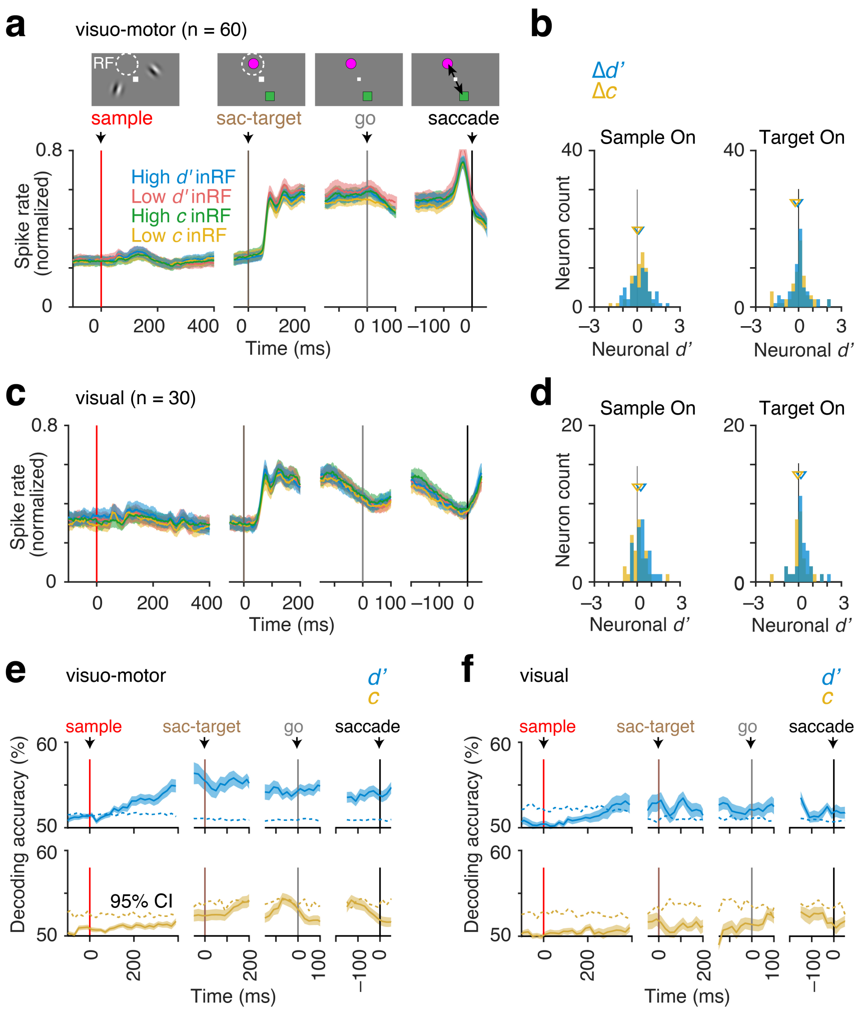


**Figure S5. Neuronal modulation in SC with behavioral $\Delta d'$ and perceptual decision $\Delta c$, when the one of the saccade targets (randomized) was placed in the neuron's RF.** (**a**) Population PSTHs of visuo-motor SC neurons when monkeys' spatial attention and decision criteria were independently controlled, and one of the saccade targets was placed at the neuron's RF location (n = 60/165). The spike rates are aligned to the onsets of sample stimuli, saccade targets, go-cue and the saccade. Error bars, ± 1 SEM. (**b**) Neuronal modulation as measured by neuronal $d'$ based on spike-counts over 200 ms (50-250 ms from the onset of sample stimuli or saccade target), associated with changes in behavioral $d'$ ($\Delta d'$) and decision criteria ($\Delta c$). (**c-d**) Similar to (a) and (b), for visual neurons. (**e-f**) Decoding accuracies for $\Delta d'$ (*top*) and $\Delta c$ (*bottom*) based on SVM linear classifiers, averaged across individual visuo-motor (e) and visual (f) neurons. Dashed lines, 95% confidence intervals based on shuffled trials. Error bars, ± 1 SEM.
